# Sound offset responses become highly informative in the auditory cortex

**DOI:** 10.1101/2025.05.19.654889

**Authors:** Charly Lamothe, Sophie Bagur, Etienne Gosselin, Brice Bathellier

## Abstract

The entire auditory system downstream of the cochlea features pronounced offset responses, which follow the termination of sounds. Because of their ubiquity, it is still an unsolved question whether offset responses are generated early in the auditory system and then propagated or recomputed at each processing stage. Here, we analysed large-scale sound responses datasets acquired in the cochlear nucleus, inferior colliculus, medial geniculate nucleus and auditory cortex of awake mice. All brain regions showed a significant proportion of offset responses often combined with onset and sustained responses in the same neuron. However, using population activity decoders, we observed that neural representations after the sound offset show a three-fold increase in sound encoding accuracy in the cortex relative to subcortical areas. This result indicates that cortical offsets encode a more precise short-term memory of the elapsed sound than subcortical offsets and that they likely result from specific computational steps.

**Key points summary:**

- Offset responses are found throughout the central auditory system
- Offset responses are often combined to sustained and onset responses at all central auditory system stages
- Offset responses provide richer information about elapsed sounds in the auditory cortex

## Introduction

Sensory processing is a dynamic phenomenon with a high sensitivity to temporal variations of the stimulus at different time scales. This is particularly acute in the auditory system, which processes complex temporal signals to extract a large range of meaningful features. Response dynamics in the auditory system include salient transient responses at the beginning and at the termination of sounds, often termed sound onset and offset respectively (Eggermont, 2015; Kopp-Scheinpflug et al., 2018). Transient onset responses are already present in the auditory nerve, where they mainly reflect adaptation, and are then observed throughout the auditory system. By contrast, no strong elevation of the firing rate is observed in the auditory nerve at sound termination, with some reports of weak inhibitory and excitatory offset responses (Grinnell, 1973; Scheidt et al., 2010; Westerman and Smith, 1984; Yin et al., 2019). Excitatory offset responses actually emerge in the cochlear nucleus (Kopp-Scheinpflug et al., 2018; Suga, 1964) and are present in all downstream stages of the main auditory system. Offset responses are thought to be important to signal sound terminations (Anderson and Linden, 2016; Li et al., 2021; Solyga and Barkat, 2021), gap detection (Awwad et al., 2020), or more generally any abrupt decrease of the sound intensity, which are part of the amplitude modulation features that the brain uses to interpret natural sounds (Bregman, 1994; Eggermont, 2015; Kuwada and Batra, 1999). Offset responses are also recruited by predictive processes, at least in the cortex, to signal omissions of expected gaps (Awwad et al., 2023). However, beyond this widely accepted but very general concept, it remains unclear what information is actually contained in offset responses and if this information is equivalent at all stages of the auditory system. On the one hand, offset responses could be an unspecific termination signal, independent of the sound that has just ended. On the other hand, offset responses could be a highly specific signal that reflects not only the termination but also the identity of the sound. For pure tones, frequency specificity of offset responses has long been established at diverse stages (Kasai et al., 2012; Kopp-Scheinpflug et al., 2018; Xie et al., 2007). This is however insufficient to assess if the offset response precisely encodes the identity of the elapsed sound, because this identity depends on complex combinations of frequency and temporal features which cannot be probed with pure tones. In order to determine the level of acoustic information in offset responses, we have analysed sound-driven activity from several thousands of neurons in two recent datasets produced by our laboratory and freely available online. These two datasets contain neural recordings at the four stages of the auditory system from the cochlear nucleus to the auditory cortex. We observed that the offset responses present at all considered stages have similar properties at the single cell level. However, strikingly, when decoding the identity of elapsed sounds based on neural population responses built from offset responses, we observed a much better performance in the auditory cortex than in subcortical regions. This marked difference suggests that cortical processing not only signals sound termination but also constructs at sound offset a rich, transiently maintained representation of the elapsed sound.

## Results

### Offset responses are prominent at all stages of the auditory system

This study is based on two datasets shared in open source repositories (Bagur and Bathellier, 2024; Gosselin, 2024). The first dataset (**Fig. 1A&B**) (Gosselin, 2024) is more extensively described in a parallel study (Gosselin et al., 2025), which does not focus on offset responses. It includes Neuropixels recordings in awake mice that sampled 1421 putative neurons (single units) in the cochlear nucleus (CN, ephys 1) and 551 single units in the inferior colliculus (IC, ephys 1). This dataset also includes two-photon calcium imaging of 4217 neurons in the auditory cortex of awake mice (AC, 2P 2). Note that response variability was larger in the auditory cortex dataset, as measured through the correlation across single trials responses for the same sound averaged across all neurons and sounds (**Fig. 1C**). For all sampled neurons of this dataset, we collected responses to 307 different sounds, each repeated 12 times. For this study, we kept 119 sounds that are all 500 ms long and contain different spectral (pure tones, frequency chords, white noise) and temporal (frequency sweeps, rhythmic and oriented amplitude modulations) features (**Fig. 1D**). The second dataset (**Fig. 1A&B**) (Bagur and Bathellier, 2024) was already used to describe population representations of simple time-varying sounds (Bagur et al., 2025). It includes silicon probe recordings in awake mice that sampled 442 putative neurons (single units) in the inferior colliculus (IC, ephys 2) and 484 single units in the medial geniculate nucleus (TH, ephys 2). This dataset also includes two-photon calcium imaging of 5936 neurons in the dorsal inferior colliculus (IC, 2P 2) and 19 414 neurons in the auditory cortex of awake mice (AC, 2P 2). As in dataset 1, larger response variability is observed in two-photon imaging data of dataset 2 as reported in (Bagur et al., 2025) with the same measure as in **Fig. 1C**. For all sampled neurons of this dataset, we collected responses to 140 different sounds, each repeated 15 times. For this study, we kept 112 sounds that are all 500 ms long and contain different spectral (pure tones, frequency chords) and temporal (frequency sweeps and oriented amplitude modulations) features (**Fig. 1D**). For extracellular electrophysiology data, single units were identified with Kilosort 2.5 (Pachitariu et al., 2024) combined with manual curation in Phy. For calcium imaging data, motion correction and neuron segmentation was performed with freely available tools (see Methods), and the data was temporally deconvolved using a simple linear algorithm allowing to achieve a temporal precision of about 3 Hz (Bagur et al., 2025) which is sufficient to separately identify onset and offset responses for the 500 ms sounds selected in this study (**Fig. 2**).

**Figure 1:**
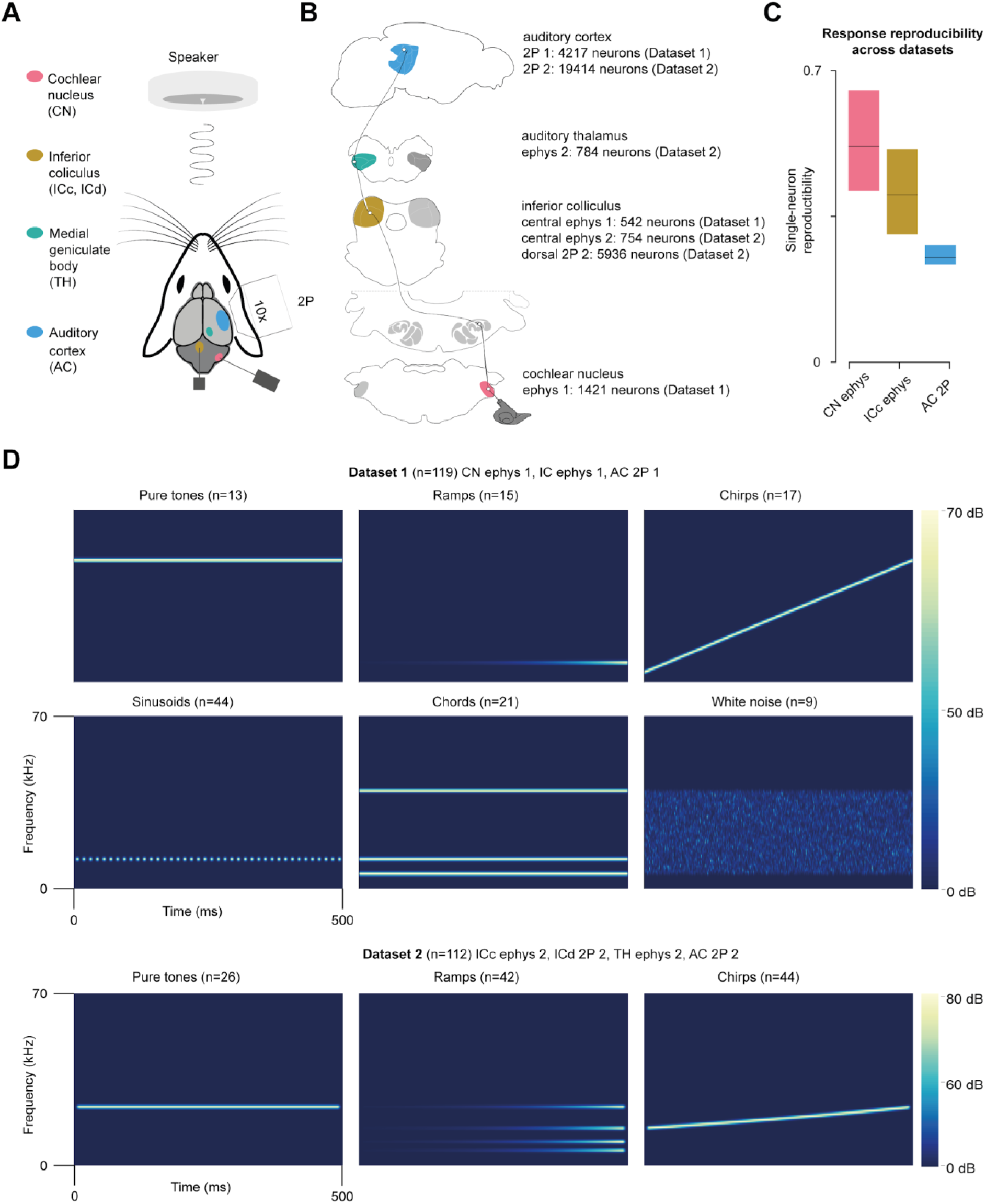
Description of the two multi-area auditory response datasets. **A.** Schematic summarizing the recording methods (extracellular multi-electrode electrophysiology: “ephys”, and two-photon calcium imaging: “2P”) and the four recorded areas. **B.** Summary of the number of putative neurons recorded in each region for each dataset. **C.** Response reproducibility across datasets. **D.** Sample spectrograms for each sound type played in each of the two datasets.

**Figure 2:**
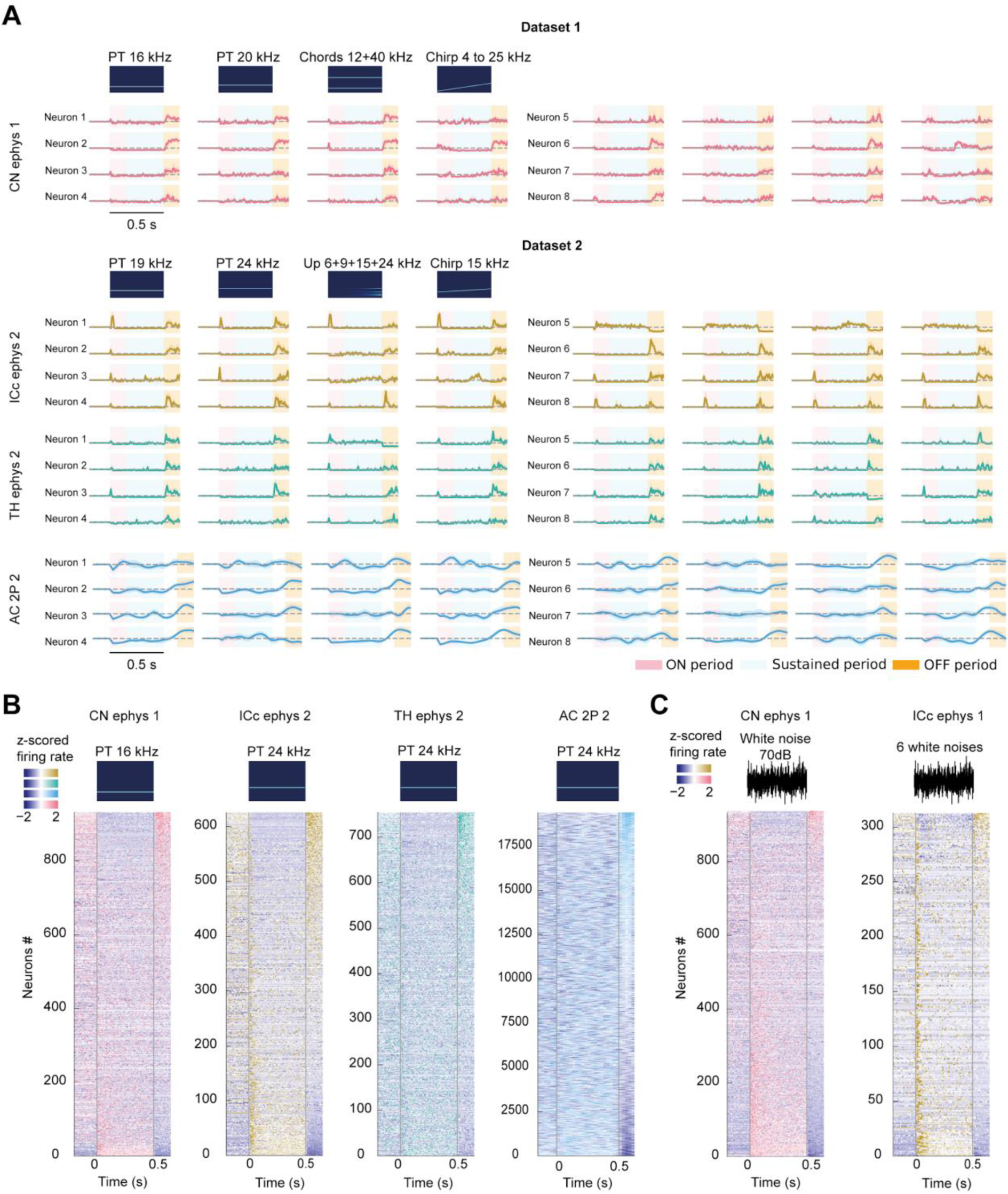
Offset responses are prominent at all stages of the auditory system. **A.** Sample responses of three selected neurons in each of the recorded areas for six different sounds. Neurons of the cochlear nucleus (CN) stem from dataset 1 and neurons of the inferior colliculus (IC), medial geniculate nucleus (TH) and auditory cortex (AC) stem from dataset 2. **B.** Heat map of the responses of all recorded neurons for the areas and datasets shown in B for one sound (Pure tone at 16 kHz/70 dB SPL and 24 kHz/80 dB SPL). **C.** (left) Heat map of the responses of responsive neurons to white noise at 70 dB SPL in the cochlear nucleus (CN ephys 1). (right) Heat map of the responses of responsive neurons to 6 different white noise sounds (different intensity profiles) in the inferior colliculus (IC ephys 1).

Plotting response traces of sample neurons from all four recorded regions, we observed, consistent with previous observations, that offset responses were present at all stages of the central auditory system. Of note, the temporal deconvolution of calcium imaging signals in the two-photon datasets (cortex and colliculus) removes the effects of the slow decay time constant of calcium dynamics but does not cancel the effects related to the slow rise time constant of GCAMP6s (∼50 to 100 ms). Therefore, the peak of offset responses in calcium imaging datasets (e.g. **Fig. 2A**) occurred typically 100 ms later than for electrophysiology recordings (**Fig. 2A**). At all stages, offset response duration could be short (few tens of millisecond) or prolonged (>100 ms), indicating that long time-scales of activity are not specific to the cortex or to calcium imaging data. This was evident both on single cell traces (**Fig. 2A**) and on heat maps representing all cell responses ordered according to the offset amplitudes (**Fig. 2B**). In the latter plots, it is also clear that offset responses can be excitatory or inhibitory (**Fig. 2B**).

Offset responses were observed for pure tones (**Fig. 2A-B**) but also for broad frequency sounds such as white noise (**Fig. 2C**). We also observed, consistent with various reports (Scholl et al., 2010; Sollini et al., 2018; Xie et al., 2007), that offset responses often co-occurred with excitatory or inhibitory onset or sustained responses (**Fig. 2A**). This indicated, as previously seen (Kopp-Scheinpflug et al., 2018), that many neurons are not dedicated to offset responses.

### Offset responses are complex at all stages of the auditory system

To evaluate if this mixed specificity of neurons to onset, sustained, or offset phases appears early or late in the auditory system, we systematically identified statistically significant responses in these three phases (**Fig. 3A**, time bins: baseline -200 to 0 ms, onset 0 to 150 ms, sustained 150 to 500 ms, offsets 500 to 650 ms for electrophysiology and 600 to 750 ms for 2P-imaging). Significant onset, sustained, and offset responses were identified by comparing firing rates during the onset and baseline periods using a non-parametric test (**Fig. 3A**, see Methods). For offset responses, we used two non-independent non-parametric tests (**Fig. 3A**) to determine if offset activity is different from both the baseline and the sustained response. For onset, sustained and offset the maximal false discovery rate was 5%, based on the alpha value of the tests (note that the two tests of offset detection being non-independent, the overall, conservative false discovery rate is also 5%, see Methods).

**Figure 3:**
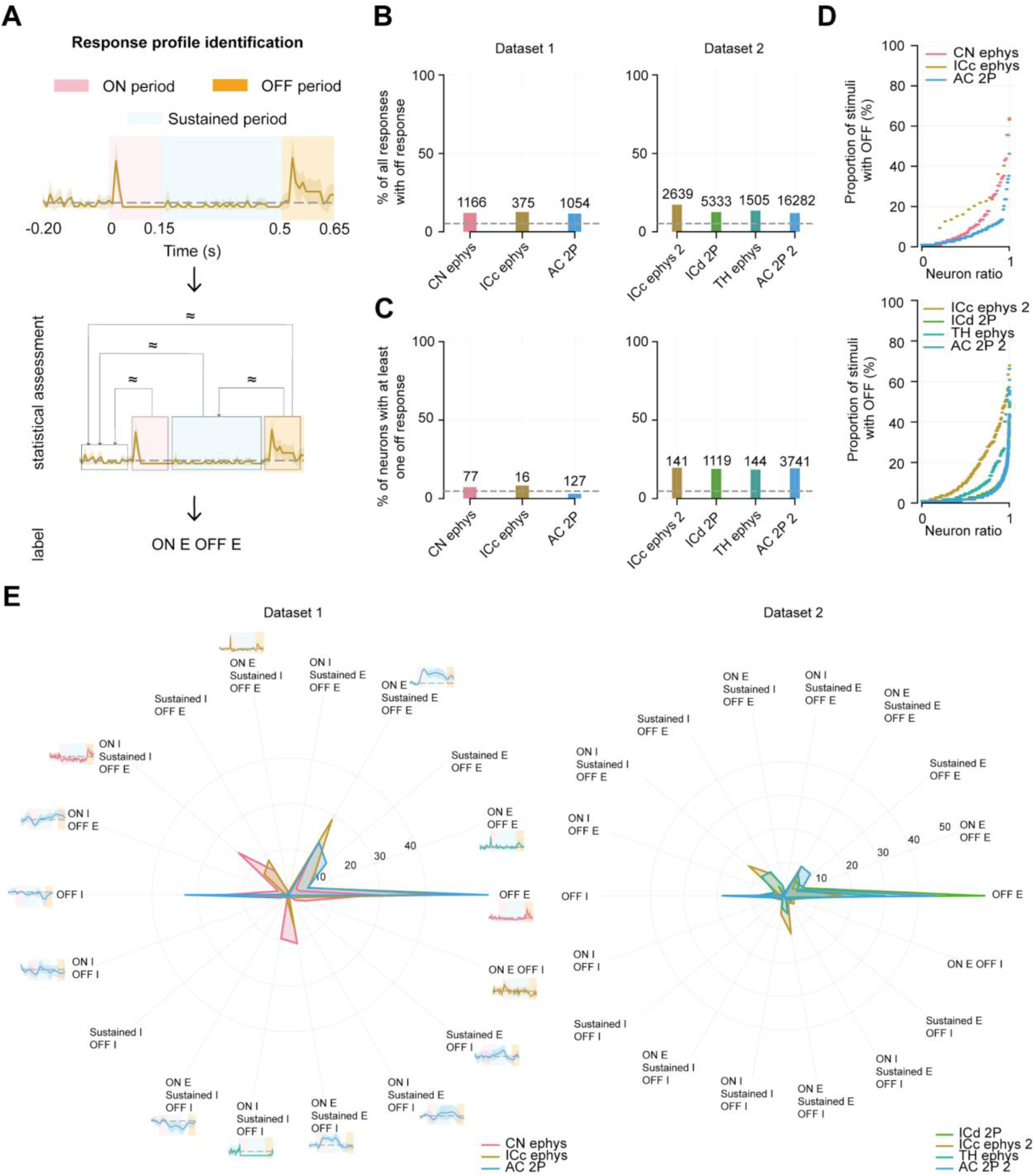
Offset responses are complex at all stages of the auditory system. **A.** Schematic of the statistical method used to detect onset, sustained and offset responses. **B.** Fraction of all pairs of sound response and neuron displaying a significant change of firing rate at the offset. Grey dotted line indicates the chance level (FDR, alpha of 0.05). **C.** Fraction of neurons for which at least one significant positive or negative offset response was detected among all tested sounds. Grey dotted line indicates the chance level (FDR, alpha of 0.05). **D.** Distribution of the fraction of responses with a significant change of firing rate (positive or negative) at the offset for each dataset. **E.** Fraction of all pairs of sound response and neuron corresponding to each possible combination of positive (E) or negative (I) onset (ON), sustained and offset (OFF) responses (18 combinations).

Based on this approach, we first quantified the fraction of all sound responses (number of sounds ✕ number of neurons) that included a significant offset response (positive or negative) at each recorded auditory system stage (**Fig. 3B**). We found this fraction to be between 11.67% and 12.65% in dataset 1 and between 11.91% and 17.31% in dataset 2, therefore systematically above the 5% chance level, consistent with the reported sparseness of offset responses. We then estimated the fraction of neurons with at least one offset response using the signed-rank tests for responses to individual sounds, but applying the Benjamini-Hochberg procedure to correct for multiple testing (119 and 112 tests per neuron in datasets 1 and 2 respectively). These estimates are likely conservative due to the low number of sound repetitions, which limits the statistical power of this offset-neuron identification procedure, with presumably many false negative detections. In dataset 1, in which we observed fewer responses (**Fig. 3B**), the fraction of neurons with detected offset is roughly equal to or below the chance level: 7.28% for CN, 8.29% for IC, 3.01% for AC (**Fig. 3C**). In dataset 2, the fraction of neurons with detected offset is clearly above chance level: 18.85% for IC_2P, 19.61% for IC_ephys, 18.44% for TH, 19.27% for AC (**Fig. 3C**). Using the results of the same statistical procedure, including correction for multiple testing, we also plotted the distribution of the number of offset responses per neuron. We observed that less than 50% of the neurons responded at the offset of more than 20 sounds, yet with an even narrower distribution for datasets derived from two-photon calcium imaging (IC and AC, **Fig. 3D**). This analysis thus suggests that single cell level measurements can be biased by the properties of the recorded signals, with, for example, an apparent smearing of offset response distributions in calcium imaging data, which is likely due to the larger signal variability with this recording modality as measured **Fig. 1C** and (Bagur et al., 2025). Beyond these discrepancies, this analysis highlights no clear progression of the offset distribution among neurons, except that IC seems to be the structure with the largest fraction of neurons displaying many offset responses compared to CN and TH (**Fig. 3D**).

We then reasoned that different auditory system stages could have different degrees of offset specificity with respect to onset and sustained responses. We therefore plotted in **Fig. 3E** the fractions of all possible combinations of an offset response (either positive, OFF E, or negative, OFF I) with onset (ON E, ON I or none) and sustained (E, I or none) responses. Note that this analysis is done response-by-response and not neuron-by-neuron, and that the fractions are given with respect to the total number of responses with a significant offset response. We observed a fraction of about 40 to 50% pure positive offset responses (OFF E) in the two-photon imaging datasets (**Fig. 3E**), while in the electrophysiology datasets, this fraction was down to 20 to 40%, with variable results across datasets (e.g. IC ephys - dataset 1: 34.58% and dataset 2: 16.97%, **Fig. 3E**). Consistent with our sample data plots of **Fig. 2B&C**, the fraction of pure negative offset was non-negligible (∼ 10 to 25% across datasets). The most prominent responses combined with excitatory offsets were sustained excitatory or inhibitory responses (Sustained E OFF E or Sustained I OFF E, **Fig. 3E**), or excitatory responses both in the onset and sustained phases (ON E Sustained E OFF E, **Fig. 3E**) or inhibitory responses both in the onset and sustained phases (ON I Sustained I OFF E, **Fig. 3E**). However, inhibitory responses before the offset were only detected in electrophysiology datasets (**Fig. 3E**). The most prominent responses combined with inhibitory offsets are monophasic onset and sustained responses, either inhibitory or excitatory (ON I Sustained I OFF E, ON E Sustained E OFF I, **Fig. 3E**). However these combinations were also only detected in electrophysiology data, and yet again with some variability, across the IC replicate which could be due to differences in stimulus properties, recording conditions, or neural population sampling (**Fig. 3E**). Altogether, this analysis of combined response type distributions highlighted clear biases across recording modalities likely related to the higher variability of calcium responses, and to their lesser sensitivity to inhibitory response (but note that many inhibitory offset responses were detected with calcium imaging). Beyond this pertaining issue, the analysis shows that, across the auditory system, offsets are often combined with earlier responses. Moreover, at all stages, there was a large fraction of responses with an excitatory offset only. Hence, we did not observe any clear evolution of neurons with offset specific responses and no clear evolution of their combination with onset and sustained response, although these differences may have been missed due to the unreliability of response detection. Both due to the rich and distributed nature of offset responses and to the difficulties encountered with signal variability at the single cell level, we reasoned that a population level analysis would be better suited to identify changes in the neural code carried by offset responses.

### Populations of offset responses are more informative in the auditory cortex

We therefore decided to measure the amount of sound information carried by the entire neural populations of each auditory system stage sampled in our dataset. Using the same analysis windows as previously defined, we constructed virtual population vectors by concatenating responses measured in all neurons across all recordings (**Fig. 4A**). Population vectors were then fed to a 5-fold cross-validated logistic regression decoder to evaluate the fraction of correct sound identity predictions that can be obtained from single trial activity of the neural population. We first established a ceiling of the information available in the full spatio-temporal activity pattern. We sampled the full time-course of the population into 65 time bins of 10 ms duration, concatenated the 65 vectors and submitted them to the same decoder. This showed similar information levels across stages, as shown previously, for dataset 2 (Bagur et al., 2025) and lower information level in cortex in dataset 1 (**Fig. 4B**), likely due to the poorer SNR in calcium imaging for dataset 1 (**Fig. 1C**). We then ran population decoders based on the time-averaged vectors obtained separately for onset, sustained, and offset time windows using the previously defined windows for each auditory system stage. To equalize across different dataset qualities, we reported the decoder accuracy computed for a specific time-bin divided by the ceiling accuracies shown in **Fig. 4B**. This analysis showed that all stages had a similar level of accuracy, with slight variations, in the onset and sustained phases of the response (**Fig. 4C**) but that accuracy was at least three fold larger in the auditory cortex than in subcortical areas in the OFF phase (**Fig. 4C**, dataset 1,CN ephys = 13.27 % +/- 1.84%, ICc ephys = 11.20 % +/- 2.99%, AC 2P = 39.60 % +/- 2.67 %, dataset 2, ICd 2P = 29.55 % +/- 2.21%, ICc ephys = 28.85 % +/- 3.99%, TH ephys = 26.91 % +/- 2.24 %, AC = 82.29 % +/- 0.63 %, p < 0.05 across all pairwise comparisons with cortex, Mann-Whitney U test, n=5 cross-validation folds).

**Figure 4:**
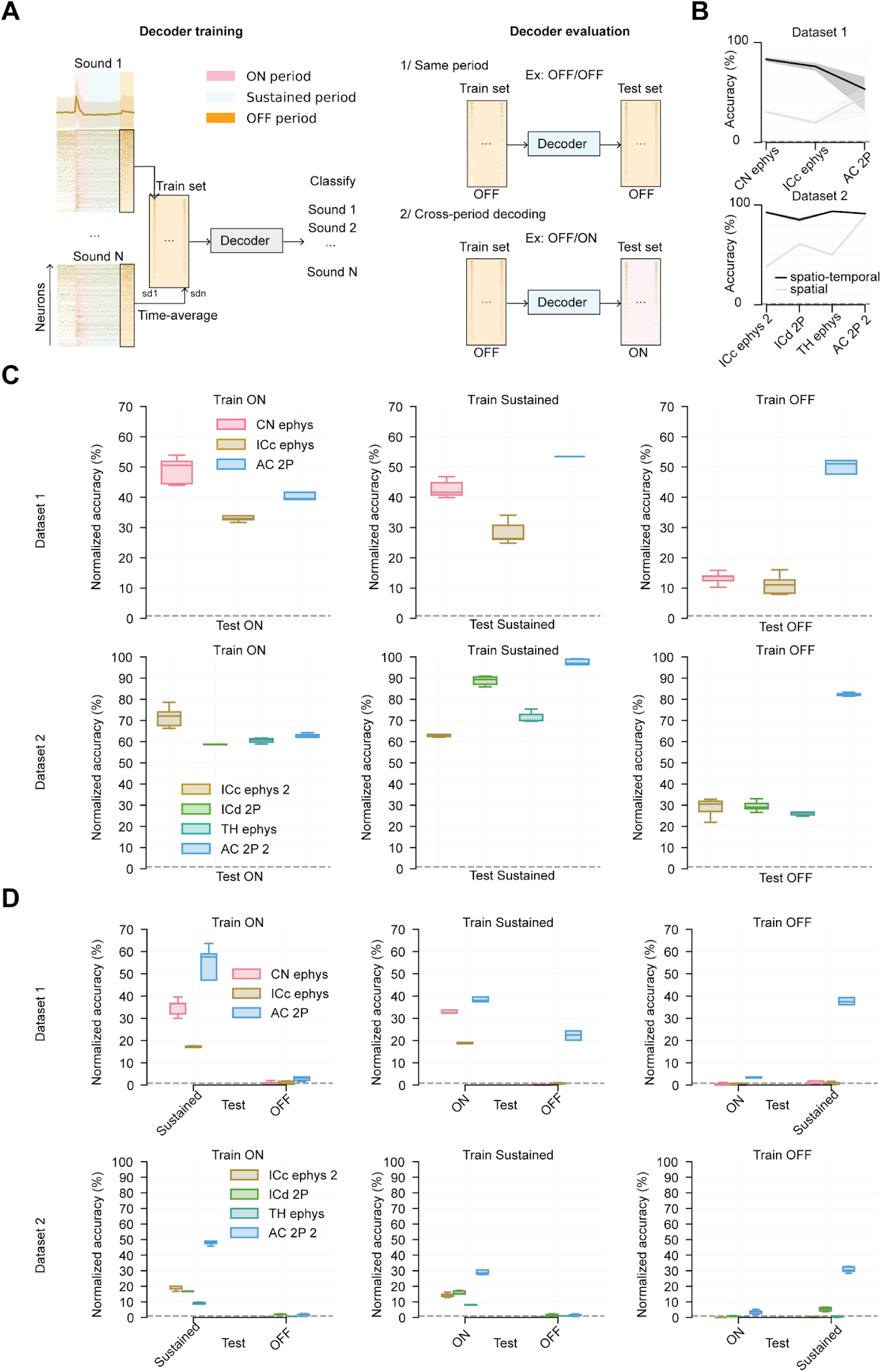
Offset responses are more informative in the auditory cortex. **A.** Schematic of the sound decoders used to predict sound identity based on neural activity. **B.** Accuracy of population activity decoders using the full spatio-temporal activity patterns (black) or only the time-averaged spatial pattern (grey). The accuracy of spatio-temporal decoders is used in C&D as the information ceiling with which the time-restricted decoder accuracy is normalized. Grey dotted line indicates the chance level (1/N sounds). **C.** Normalized accuracy of population decoders based on time-averaged firing rates computed during the onset, sustained and offset phases of the responses respectively for all datasets and recorded brain regions. The normalized accuracy represents the fraction of the ceiling accuracy estimated with the full spatio-temporal decoder (see B). Grey dotted line indicates the chance level. **D.** Normalized accuracies for decoders trained in one response phase but tested on another response phase. Grey dotted line indicates the chance level.

Both in datasets 1 and 2, cortical data are derived only from calcium imaging, raising the possibility that the very important difference of decoder performance could be due to spurious effects of this recording modality. Specifically, temporally deconvoled calcium imaging data are still temporally smeared by the slow rise time of GCAMP6s which is visible for example in the delayed peak of offset responses (**Fig. 2A**). We have compensated for this effect by delaying the offset analysis window for calcium data (**Fig. 3**). Nevertheless, it is difficult to rule out *a priori* a transfer of sustained response information into the offset window in calcium imaging data due to this temporal smearing. Three observations allow us to exclude this possibility. First, decoder accuracy was poor for inferior colliculus offsets both for electrophysiology and calcium imaging data (**Fig. 4C**). Second, decoders trained on auditory cortex sustained responses had poor accuracy when tested on cortical offset responses (**Fig. 4D**, AC 2P 1: 22.21 % +/- 1.82 %, AC 2P 2: 1.47 % +/- 0.54 %), suggesting that the information used by the decoder during the offset does not simply mirror sustained activity. However, decoders of offset responses displayed an intermediate accuracy level when tested on sustained activity (**Fig. 4D**, AC 2P 1: 33.68 % +/- 2.30 %, AC 2P 2: 30.71 % +/- 1.67 %) suggesting a degree of overlap between sustained and offset responses. This can have multiple causes including temporal smearing, but also a partially shared representation space between sustained and offset phases in AC.

Therefore, our third observation aimed at discriminating between these two alternatives. We decomposed the calcium response on each trial into a linear sum of a set of onset, sustained and offset regressors which modeled the temporal smear based on our data (**Fig. 5A**). Most importantly, we modeled sustained response with a decaying component that extended into the offset time window (**Fig. 5A**). The regressors have a fixed time course but three time-dilated regressors were used at the offset to capture the variability of response time constants across neurons (**Fig. 5A**). This approach allowed us to extract one amplitude value for onset, sustained and offset responses while canceling largely the smearing effect. We verified the latter aspect by plotting the amplitude of sustained against offset components and comparing this with the relation of time-averaged values during sustained and offset periods for calcium imaging data. A clear drop of the correlation between offset and sustained with the linear demixing model confirmed its efficiency to cancel the temporal smearing effect in calcium signals (**Fig. 5B**). We then measured the accuracy of the decoders trained on the offset component and tested on the sustained component of the demixing model. We observed overall only a modest drop in classification accuracy compared to response averaging, indicating that the smearing effect, largely canceled by demixing, contributes little to the information contained in offset responses in the auditory cortex dataset (**Fig. 5C**). Likewise, the cross-decoding of sustained activity by offset decoders was little impacted by the demixing model, showing that temporal smearing does not explain this effect, which thus rather corresponds to a genuine overlap between sustained and offset representation spaces in the cortex (**Fig. 5D**).

**Figure 5:**
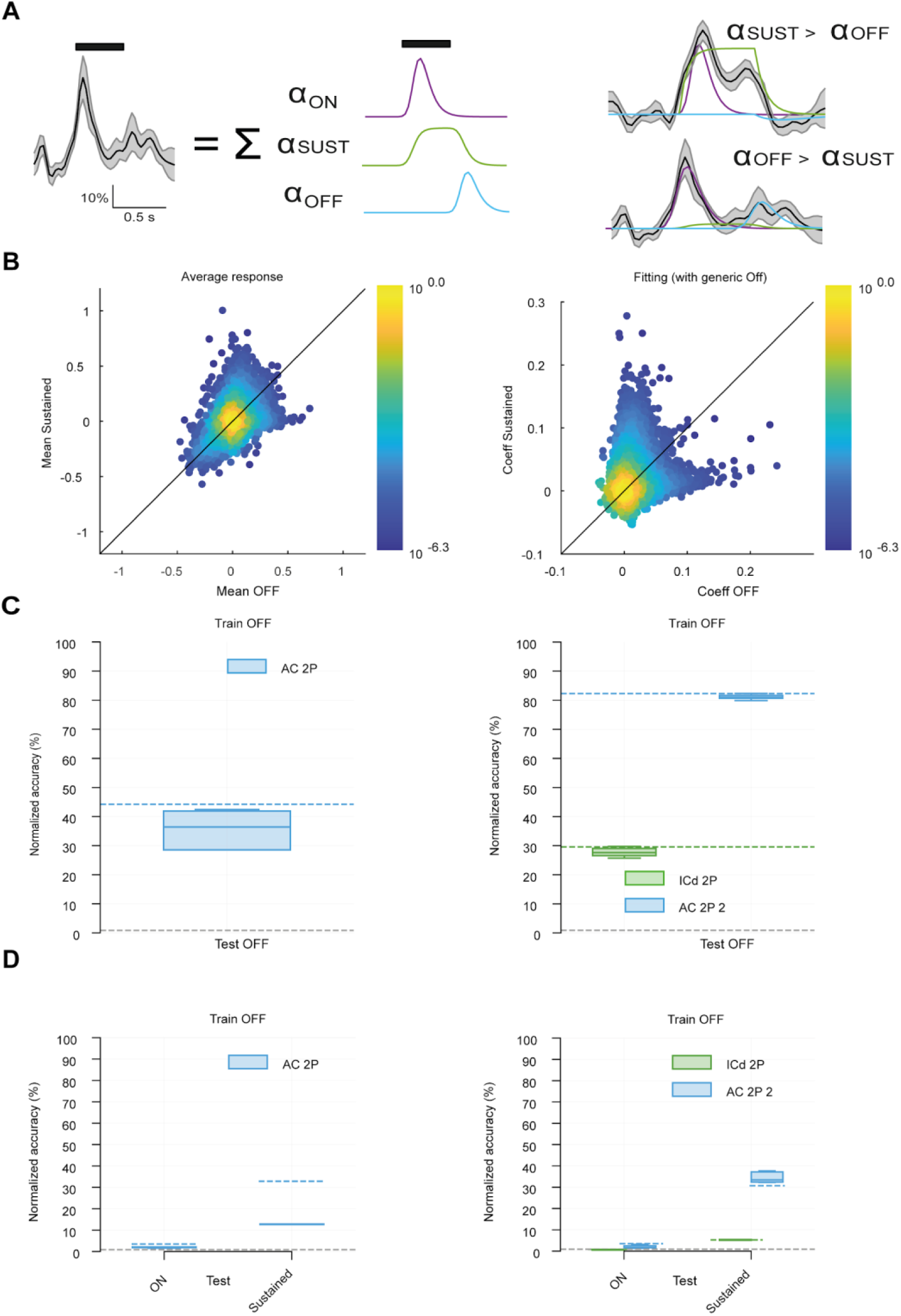
Offset information is specific to offset responses. **A.** Schematic of the response demixing algorithm used to separate smeared sustained responses from real offset responses in two-photon calcium imaging. **B.** (left) Scatter plot of all the time averaged deconvolved calcium signals in the offset and sustained time window showing a strong correlation between the two values. (right) Scatter plot of all the fitted offset and sustained coefficients for the demixing models in the offset and sustained time window showing a decorrelation of the offset and sustained component of sound responses. **C.** Decoder accuracies based only on the offset component of the response model compared to accuracies based on raw data (blue and green dotted lines identical to Fig. 4). Grey dotted line indicates the chance level. **D.** Accuracy of the decoder based on the offset component of the response model when tested on the onset and sustained components of the response model. Grey dotted line indicates the chance level.

## Discussion

Our results provide a systematic analysis of offset responses across the auditory system based on two large datasets collected in awake mice. They complement a wide range of studies that have reported the presence of offset responses throughout the auditory system and characterized their properties across auditory system stages and animal species (Kopp-Scheinpflug et al., 2018). The main specificities of our dataset are the large number of neurons recorded especially with calcium imaging and the homogeneity of the stimulus set and recording conditions across the different stages. Also, some recordings (e.g. inferior colliculus or auditory cortex present in dataset 1 and 2, calcium and electrophysiology in inferior colliculus) provide repeats of similar experiments allowing to verify the stability of observed properties across recording modalities, experimental conditions and sampling biases. Our datasets represent a unique collection of mouse auditory system data, but are incomplete to some degree with the absence of recordings in the superior olivary complex and the lack of thalamic data in dataset 1 or of cochlear nucleus in dataset 2.

Reassuringly, our broad dataset recapitulates the main observations made in the past about offset responses. First, offset responses are found at all stages (**Fig. 3**) in a small subset of neurons. Although consistent with observations that offset responses are sparse, our measurement of the fraction of neurons with offset responses may underestimate the real value due to the poor statistical power of our datasets for this type of question (low sound repeat number and large number of sounds). Previous studies with more statistical power indeed reported larger fractions of offset responsive neurons in the inferior colliculus (Akimov et al., 2017) or in the auditory cortex (Tian et al., 2013). In the cochlear nucleus, offset responses are thought to be present only in specific subtypes of neurons (Ding et al., 1999; Young and Brownell, 1976). It is possible that the large representation of offset neurons in our cochlear nucleus dataset is due to a sampling bias introduced by Neuropixels recordings.

Second, offset responses are often, although not systematically, combined with onset and sustained responses (**Fig. 3**), a property commonly reported at different stages of the auditory system (Kopp-Scheinpflug et al., 2018). Interestingly, the frequency tuning of on- and offset responses in the same neurons of the auditory cortex is often different (Scholl et al., 2010; Sollini et al., 2018) suggesting that on- and offset responses in cortex come from circuit processes that are more complex than a single, shared input channel.

Our population decoding analysis demonstrates that cortical offset responses are not only distinct from onset responses but they are also much more informative about the identity of the elapsed sound than subcortical offsets (**Figs. 4-5**). The classifier accuracy improvement between cortical and subcortical levels is more than 3-fold in our two datasets, indicating a robust and clear effect. This striking difference is not due to the large sample size of auditory cortex recordings. Two-photon calcium imaging data are noisier (**Fig. 1C**) (Bagur et al., 2025) and display sparser offset responses than electrophysiology data (**Fig. 3D**). These two biases are compensated by the larger sample size of the two-photon calcium imaging data. This is shown in our ceiling information analysis (**Fig. 4B**), which uses population activity decoders based on the full spatio-temporal information available in each neuron to estimate the maximum sound information available in the dataset. This analysis shows that ceiling information is similar across datasets (**Fig. 4B**). Moreover, decoder accuracies are more similar across auditory system stages for onset and sustained responses (**Fig. 4B**). These two observations demonstrate that the larger sample size of two-photon calcium imaging does not provide additional information which could explain the highly informative offset in AC. Also, if the sample size of two-photon calcium imaging data would be a factor, offset responses should also be more informative in the two-photon calcium imaging dataset than in the electrophysiology dataset recorded in the inferior colliculus, which we do not observe (**Fig. 4C**). The boost of offset information is also not due to a spill over of sustained information in cortical recordings as demonstrated with our linear demixing analysis (**Fig. 5**). Hence, we conclude that auditory cortex response dynamics are such that they generate highly informative offset activity that does not exist upstream.

This observation is a clear indication that offset responses do not result from a combination of labelled-line offset inputs from upstream areas. Indeed, if this were the case, as information cannot be created *de novo*, the decoder accuracy would be as poor in the cortex as subcortically. Our results therefore show that offset responses in the auditory cortex are recomputed not only from upstream offsets but also necessarily using information originating in earlier phases of the response, which are much more informative than offset responses at subcortical levels. A previous study reported an absence of offset responses to white noise in the thalamus but not in the auditory cortex (Solyga and Barkat, 2021), while offset responses were present in both stages for pure tones. This suggested that cortical offset includes new information as compared to subcortical offsets. Our data do not corroborate this observation, as we observe clear offset responses to white noise even in the inferior colliculus (**Fig. 2C**). Our observations therefore provide a strong argument for the idea of newly computed offset responses in the auditory cortex.

What processes could explain our observation of richer offset responses in the auditory cortex? Improved decoder performance in the cortex could result from either an improvement of sound-specific signals or a decrease of offset response variability or both. As response variability was larger in cortical datasets, it is likely that decoder performance was rather improved by an increased sound-specificity of cortical offsets compared to their subcortical counterparts. The mechanisms usually proposed for bottom-up generation of offset responses divided between rebound mechanisms in single neurons and local micro-circuits mechanisms providing for example time-shifted inhibitory and excitatory inputs (Kopp-Scheinpflug et al., 2018). Because of their spatial locality, these mechanisms are not sufficient to identify the factors underlying the richness of offset responses in complex circuits with multiple inputs. Recently, it was demonstrated that large recurrent neural networks produce complex offset responses that depend on multiple interactions within the network (Bondanelli et al., 2021). Anatomically, all auditory system networks considered in this study have a recurrent inhibitory and excitatory architecture (Levy and Reyes, 2012; Oliver, 2005; Young and Davis, 2002). It is therefore conceivable that offset responses originate from recurrent network-scale interactions. This hypothesis would generalize local micro-circuit models. It would be highly compatible with the high sensitivity of offset responses to anesthetics that strongly affect network dynamics (Filipchuk et al., 2022; Gosselin et al., 2024). In this framework, our results indicate that the collective dynamics generating offset responses are more complex in the auditory cortex than in upstream stages. This complexity could span both the spatial dimension (i.e. the extent to which interactions involve multiple neurons processing multiple sound features) and the temporal dimension (i.e. the temporal span over which information is integrated to produce the offset response). It is already well established that auditory cortex responses have less temporal precision than subcortical responses (Wang et al., 2008) and are influenced by a longer history time-window (Norman-Haignere et al., 2022). This could contribute to the higher richness of cortical offset responses, which due to their long duration (Kopp-Scheinpflug et al., 2018) (**Fig. 2**) also provide, at the population scale, a transient memory of the elapsed sound.

## Material availability

Biological material and technologies used in this study are freely available resources.

## Data and code availability

All datasets and custom codes will be freely available on Zenodo.

## Acknowledgments

We acknowledge the support of the Fondation pour l’Audition to the Institut de l’Audition.

We acknowledge the support of the following funding sources: Fondation pour l’Audition, RD-2023-1 (BB), FPA IDA02 (BB) and APA 2016-03 (BB), European Research Council, ERC CoG 770841 DEEPEN (BB) Fondation pour la Recherche Médicale SPF202005011970 (SB) Agence Nationale pour la Recherche, France 2030 program, ANR-23-IAHU-0003 (BB)

## Author contributions

Conceptualization: CL, EG, BB, SB

Methodology: CL, EG, BB, SB

Investigation: CL, EG, SB

Data curation: CL, EG, SB

Formal analysis: CL, EG, SB

Supervision: BB, SB

Project administration: BB

Funding acquisition: BB, SB

Software: CL, SB

Visualization: CL

Validation: BB, SB

Writing-original draft: CL, EG, BB, SB

Writing-review & editing:

## Declaration of interests

The authors declare no competing interests.

## Declaration of generative AI and AI-assisted technologies

The authors did not use generative AI and AI-assisted technologies at any step of the study and in the writing process.

## Data availability

All data supporting the results in the paper are archived in the public repository Zenodo (access links: doi:10.5281/zenodo.13941450 and doi:10.5281/zenodo.14421103).

## Material and Methods

### Ethical approval

All experimental and surgical procedures were carried out in accordance with the European guidelines for animal experiments and were validated by the French Ethical Committee #89 (authorization APAFIS#27040-2020090316536717 v1).

### Datasets

All data used in this study are derived from two publicly available datasets summarized in **Table 1**. Dataset 2 was described more extensively in a published article (Bagur et al., 2025). Dataset 1 is described in a parallel pre-print manuscript (Gosselin et al., 2025).

**Table 1.** Summary of the datasets.

| Dataset 1 |  |  |  |  |  |  |
| --- | --- | --- | --- | --- | --- | --- |
|  | Method | N animals | N session | N neurons | N good neurons | Soundset |
| CN | Electrophysiology (1x384) | 8 | 21 | 2265 | 1421 (63%) | 119 sounds (pure tones, chords, ramps, frequency sweeps, white noise) |
| IC - Central | Electrophysiology (1x384) | 9 | 15 | 1197 | 551 (46%) |  |
| AC | 2 photon calcium imaging | 5 | 15 | 14055 | 4217 (30%) |  |

**Table 1. Summary of the datasets**
| Dataset 2 |  |  |  |  |  |  |
| --- | --- | --- | --- | --- | --- | --- |
|  | Method | N animals | N session | N neurons | N good neurons | Soundset |
| IC - Central | Electrophysiology (1x32) | 11 | 30 | 563 | 442 (78%) | 112 sounds (pure tones, up/down ramps, up/down frequency sweeps) |
| IC - Dorsal | 2 photon calcium imaging | 30 | 101 | 15312 | 5936 (39%) |  |
| TH | Electrophysiology (4x8) | 10 | 33 | 498 | 484 (97%) |  |
| AC | 2 photon calcium imaging | 7 | 60 | 60822 | 19414 (32%) |  |

### Subjects and ethical authorizations

All mice used for the production of the datasets were 10 to 16 weeks old male C57Bl6J mice (26 - 30 g) obtained from Janvier labs (Le Genest-Saint-Isle, France) that had not undergone any other procedures. Mice were group-housed (2–4 per cage) before and after surgery, had *ad libitum* access to food and water and enrichment (running wheel, cotton bedding and wooden logs) and were maintained on a 12-hour light-dark cycle in controlled humidity and temperature conditions (21-23 °C, 45-55% humidity). All experiments were performed during the light phase. The terminal procedure was carbon dioxide inhalation.

### Surgical procedures

Mice were injected with buprenorphine (Vétergesic, 0,05-0,1 mg/kg) 30 min prior to surgery. Surgical procedures were carried out using either intraperitoneal ketamine (Ketasol) and medetomidine (Domitor) which was antagonized with atipamezole (Antisedan, Orion pharma) at the end of the surgery) or 3% isoflurane delivered via a mask. After induction, mice were kept on a thermal blanket during the whole procedure and their eyes were protected with Ocrygel (TVM Lab). Lidocaine was injected under the skin of the skull 5 minutes prior to incision.

For electrophysiology recordings, the skull was exposed at the relevant location for ulterior craniotomy: above the IC, above the cortex dorsal to the TH or above the dorsal cerebellum for the CN. A well was formed around it using dental cement in order to retain saline solution during recordings and the head post was fixed to the skull using cyanolit glue and dental cement. To protect the skull, the well was filled with a waterproof silicone elastomer (Kwikcast, WPI) that could be removed prior to recording. The head post was fixed to the skull using cyanolit glue and dental cement (Ortho-Jet, Lang).

For calcium imaging, craniotomies of 3 (IC) or 5 (AC) mm were performed above the IC or the AC. Injections of 150nL of AAV1.Syn.GCaMP6s.WPRE (Vector Core, Philadelphia, PA; 10^13 viral particles per ml; used pure for TH and diluted 30x for AC and IC) were made at 30 nL/min with pulled glass pipettes at a depth of 500µm and spaced every 500 µm to cover the large surface of the IC or AC. The craniotomy was sealed with a circular glass coverslip. The coverslip and head post were fixed to the skull using cyanolit glue and dental cement (Ortho-Jet, Lang).

After surgery, mice received a subcutaneous injection of 30% glucose and metacam (1 mg/kg). Mice were subsequently housed for one week with metacam delivered via drinking water or dietgel (ClearH20). Mice were given one week to recover from surgery without any manipulation. Then, for four days before recording, mice were habituated to head restraint for increasing periods of time (30 min - 2 hours). For electrophysiological experiments, the day before recording animals were briefly anesthetized using isoflurane anesthesia (2%) in order to perform craniotomy and durectomy for electrode descent.

### Electrophysiological recordings

Dataset 1 (CN): The awake mouse was head-fixed and Neuropixels 1.0 probes (384 channels) were inserted through the cerebellum at a 38-40° angle in the sagittal plane, targeting the contralateral cochlear nucleus, vertically targeting the inferior colliculus, or both. Electrode angle and entry point were defined relative to the initial head-post placement. Fine tuning of these targeting parameters was progressively obtained through repeated penetrations based on time-locked responses to sounds easily detectable during probe insertion. Data was sampled at 30kHz using a NI-PXI chassis (National Instruments) and the SpikeGLX acquisition software.

Dataset 2 (IC, TH): The awake mouse was head-fixed and Neuronexus 1x32 linear probe (IC) or 4x8 comb prove (TH) was inserted. Data was sampled at 20kHz using an Intan RHD2000 amplifier board.

Recording was started 15-20 minutes after the electrode position was locked to allow the brain tissue to stabilize and minimize movements of neurons in the first part of the recording. Recordings were performed using warmed saline filling the cyanolit glue well and in contact with the reference electrode. After each recording the well was amply flushed and then refilled with Kwik-Cast.

For post-hoc histological verification of the electrode track using fluorescent dye, the electrodes were dipped in diI, diO or diD (Vybrant™ Multicolor Cell-Labeling Kit, Thermofisher) prior to insertion.

### Two-photon calcium imaging

Dataset 1 (AC): Imaging was performed using an acousto-optic two-photon microscope (Karthala System, Orsay, France) ewith a pulsed laser (Insight, Spectra-Physics, Santa Clara, CA) set at 900 nm. We used a 16x objective (N16XLWD-PF, Nikon) to acquire stacks of four images at 22.9 Hz from four planes interleaved by 50 µm in a field of view of 478x478 µm.

Dataset 2 (AC and IC): Imaging was performed using a two-photon microscope (Femtonics, Budapest, Hungary) equipped with an 8kHz resonant scanner combined with a pulsed laser (MaiTai-DS, SpectraPhysics, Santa Clara, CA) set at 900 nm. We used a 10x Olympus objective (XLPLN10XSVMP), which provided a field of view of up to 1x1 mm. For AC, a 1x1mm field of view was used. For IC, the field of view was adjusted to the size of the structure (∼0.5x0.5 mm). Images were acquired at 31.5 Hz.

### Sound set and experimental protocol

Dataset 1: Sounds were generated with Python (The Python Software Foundation, Wilmington, DE) and were delivered at 192 kHz with Matlab (The Mathworks, Natick, MA), using a NI-PCI-6221 card (National Instruments) driven by a custom protocol using the Matlab Data Acquisition toolbox and feeding an amplified free-field loudspeaker (SA1 and MF1-S, Tucker-Davis Technologies, Alachua, FL) positioned in front of the mouse, 10 to 15 cm from the mouse ear. Sound intensity was cosine-ramped over 10 ms at the onset and offset to avoid spectral splatter. The head fixed mouse was isolated from external noise sources by sound- proof boxes (custom-made by Decibel France, Miribel, France) providing 30 dB attenuation above 1 kHz. Sound pressure levels were computed as Root Mean Square. During a recording session, 307 sounds were repeated 12 times and played in a random order with a 1 s interval between sound onsets in 123 blocks of 30 sounds (total duration ∼80 minutes). The 307 sounds included simple and more complex sounds. Because of the strong time-varying nature of the complex sounds we focused the analysis of offset responses on a set of 152 more simple sounds whose duration was 500ms. These included 28 Pure tones: pure tones at 14 frequencies logarithmically spaced between 2 kHz and 60 kHz at 50 dB SPL and 70 dB SPL. 26 Ramps: linearly ramped sounds, increasing and decreasing in intensity, at the same frequencies as the 13 pure tones between 2 kHz and 50 kHz between 50 dB SPL and 70 dB SPL. 48 Chords: summation of 2-4 70 dB pure tones from low (11 sounds), medium (11 sounds), high (11 sounds) frequency groups, broadly sampled frequencies (10 sounds) and harmonically arranged frequencies (5 sounds). 20 Chirps: up- and down-frequency sweeps of different durations between 25 ms and 400 ms at 6-12 kHz, 50 dB SPL (10 sounds) or different frequency content between 4 kHz and 50 kHz at 50 dB SPL in 500 ms (10 sounds). 30 Coloured noises (WN): broadband noises at 50 dB SPL, 70 dB SPL or up-/down-ramped (6 sounds), filtered noises at different bandwidths between 2 kHz and 80 kHz (14 sounds), and summation of two 1 kHz bandwidth filtered noises up- and down-ramped between 50 dB SPL and 70 dB SPL, matching frequencies of a subset of chords (10 sounds). 48 amplitude-modulated sounds (AM): sinusoidally amplitude modulated sounds at 6 different modulation frequencies between 4 Hz and 160 Hz and 8 carrier frequency contents (2 pure frequencies, 5 sums of frequencies matching chords and 1 broadband noise). Out of the 152 sounds, 33 produced unreliable population responses (mean correlation across time-averaged single trial population vectors < 0.1) in at least one of the three recorded areas. These sounds were excluded from the final analysis, restricted to 119 sounds, to focus on robust responses. We verified that this had no effect on our conclusions.

Dataset 2: Sounds were generated with Matlab (The Mathworks, Natick, MA) and were delivered at 192 kHz with a NI-PCI-6221 card (National Instruments) driven by the software Elphy (G. Sadoc, UNIC, France) and feeding an amplified free-field loudspeaker (SA1 and MF1-S, Tucker-Davis Technologies, Alachua, FL) positioned 15 to 20 cm from the mouse ear. Sound intensity was cosine-ramped over 10 ms at the onset and offset to avoid spectral splatter. The head fixed mouse was isolated from external noise sources by sound-proof boxes (custom-made by Femtonics, Budapest, Hungary or Decibel France, Miribel, France) providing 30 dB attenuation above 1 kHz. Sounds were calibrated in intensity at the location of the mouse ear using a probe microphone (Bruel & Kjaer, type 4939-L-002). For two-photon calcium imaging, the resonant scanner generated a harmonic background noise at 8 kHz (intensity at the mouse ear, 45 dB SPL). During a recording session, each sound was presented 15 times in random order.

The set of sounds consisted of 112 sounds (Fig. 1) which were all 500 ms long. 26 Pure tones: pure tones at 11 frequencies logarithmically spaced between 4 kHz and 37 kHz at 50, 70 and 80 dB SPL. 42 Ramps: linearly ramped sounds, increasing and decreasing in intensity, at various frequencies and combinations (chords) of frequencies. 44 Chirps: up- and down-frequency sweeps with various start and stop frequencies between 4 and 24 kHz at 50 and 80dB. These sounds were played in pseudo random order together with 28 other sounds (shorter frequency ramps and long amplitude modulated sounds, see (Bagur et al., 2025)) which were excluded from the present analysis.

## Data preprocessing

Dataset 1: Motion artifacts, regions of interest selection, and the signal extraction were carried out with the standard pipeline of the Python-based version of Suite2p (Pachitariu et al., 2017). For each region of interest, the mean fluorescence signal *F(t)* was extracted together with the local neuropil signal *F*_np_*(t)*. Then 70% of the neuropil signal was subtracted from the neuron signal to limit neuropil contamination. Baseline fluorescence F_0_ was calculated with a sliding window computing the 3rd percentile of a Gaussian-filtered trace over the imaging blocks. Fluorescence variations were then computed as *f(t)* = Δ*F/F* = (*F(t)* - *F*_0_)/*F*_0_ . An estimate of firing rate variations r(t) was then obtained by linear temporal deconvolution of f(t): *r(t)* = *f’(t) + f(t)/τ, f’(t)* being the first derivative of *f(t)* and *τ* = 2s, the estimated decay of the GCAMP6s fluorescent transients. This simple method efficiently corrects the strong discrepancy between fluorescence and firing rate time courses due to the slow decay of spike-triggered calcium transients. It does not correct for the rise time of GCAMP6s, leading to remnant low pass filtering of the firing rate estimate and a delay of ∼100ms between the firing rate peaks and the peaks of the deconvolved signal. Finally, response traces were smoothed with a Gaussian filter (σ = 31ms). Single trial sound responses were extracted (0.2 s before up to 0.8 s after sound onset) and the average activity over the prestimulus period (0.2 s - 0.02 s before sound onset) was subtracted for each trial.

For electrophysiology, raw data were band-pass filtered (300 - 6 000 Hz) and channels from the electrode tip (corresponding to the cochlear nucleus region) were selected using SpikeInterface (https://github.com/SpikeInterface) (Buccino et al., 2020). Isolated clusters were identified using Kilosort 2.5 (Pachitariu et al., 2024) followed by manual curation based on the interspike-interval histogram and the inspection of the spike waveform using Phy (https://github.com/cortex-lab/phy). Canonical spike sorting was first applied with common parameters throughout the whole recording, attempting to optimize the spike detection and assignment to clusters. The measure of drift throughout the recording computed with Kilosort 2.5 showed minimal slow drift throughout the recording, except for some experiments in the first 10 minutes. Due to the high spiking activity in the cochlear nucleus, optimal spike sorting was achieved using lower Kilosort 2.5 parameters (detection threshold = 6, clustering threshold = 6, and matching thresholds = [9,4]) in this region than for the inferior colliculus (detection threshold = 8, clustering threshold = 8, and matching thresholds = [10,4]). After manual curation, single trial sound responses were extracted (0.2 s before up to 0.8 s after sound onset) as a histogram of 1ms time bin and the average activity over the prestimulus period (0.2 s - 0 s before sound onset) was subtracted for each trial. Based on histology, we identified a number of units whose location on the Neuropixel probe was not compatible with a localization in the cochlear nucleus or inferior colliculus.

### Dataset 2

For calcium imaging, regions of interest corresponding to putative neurons (AC 2P and IC) or axons and boutons (TH ephys) were identified by using Autocell (Deneux et al., 2016) (https://github.com/thomasdeneux/Autocell). Briefly, each frame of the recording was corrected for horizontal motion using rigid body registration. This step was visually controlled and all sessions with visible z motion were eliminated. A hierarchical clustering algorithm, based on pixel covariance over time, agglomerated pixels up to a user-selected number of clusters corresponding to regions of the size of neurons of axons. Clusters were automatically filtered according to size and shape criteria. This step was controlled by a detailed visual inspection of selected regions of interest (ROIs) during which ROIs without visually identifiable cell body shape were discarded.

For each region of interest, the mean fluorescence signal *F(t)* was extracted together with the local neuropil signal *F_np_(t)*. Then 70% of the neuropil signal was subtracted from the neuron signal to limit neuropil contamination. Baseline fluorescence F_0_ was calculated with a sliding window computing the 3rd percentile of a Gaussian-filtered trace over the imaging blocks. Fluorescence variations were then computed as *f(t) = ΔF/F = (F(t) - F_0_)/F_0_* . An estimate of firing rate variations *r(t)* was then obtained by linear temporal deconvolution of f(*t*)*: r(t) = f’(t) + f(t)/τ, f’(t)* being the first derivative of *f(t)* and *τ* = 2s, the estimated decay of the GCAMP6s fluorescent transients. This simple method efficiently corrects the strong discrepancy between fluorescence and firing rate time courses due to the slow decay of spike-triggered calcium transients. It does not correct for the rise time of GCAMP6s, leading to remnant low pass filtering of the firing rate estimate and a delay of ∼100 ms between the firing rate peaks and the peaks of the deconvolved signal. Finally, response traces were smoothed with a Gaussian filter (*σ* = 31 ms).

Electrophysiological signals were high-pass filtered and spike sorting was performed using the CortexLab suite (https://github.com/cortex-lab, UCL, London, England). Single unit clusters were identified using Kilosort 2.5 (Pachitariu et al., 2024) followed by manual corrections based on the interspike-interval histogram and the inspection of the spike waveform using Phy (https://github.com/cortex-lab/phy).

Both for imaging and electrophysiology data, single trial sound responses were extracted (0.5s before and 1s after sound onset) and the average activity over the prestimulus period (0.5s - 0s before sound onset) was subtracted for each trial.

### Reproducibility index and cell selection

To quantify the noise levels in the data, we calculated the mean inter-trial correlation across all pairs of trials. The single neuron reproducibility is then defined for each neuron as the average of the inter-trial correlation for that neuron’s response to all sounds presented in the experiment. Region of interests (ROIs) or single units with reproducibility below 0.12 were classified as non-responsive and were excluded from all analyses. The number of responsive units and the corresponding fraction of the total number of units/ROIs are given in **Table 1**.

### Population activity decoding

To decode the stimulus identity from neural activity, we evaluated performance across different response windows, including onset, sustained, offset, and full response periods. For each window, neural responses from all neurons in each considered dataset were extracted and concatenated into a virtual population vector. In order to avoid statistical threshold effects, the selection of offset responses based on single neuron responses was not used in the population decoding approach. These vectors were constructed by averaging neural responses across the analysis time bin (onset, sustained, offset) except for the measurement of ceiling information. For the latter analysis, 65 time bins were defined from 0 to 150 ms after stimulus onset (650 ms in total), a population vector was constructed for each time bin and vectors from all time bins were concatenated into a vector summarizing the full spatio-temporal activity time-course of the population.

Next, the neural responses from each window were used as input to a stratified 5-fold cross-validation logistic regression decoder. Stratified cross-validation ensured that the class distributions were preserved in each fold, addressing any class imbalances. During each fold, training and test set were normalized by subtracting the training set mean. This approach allowed us to assess the fraction of correct sound identity predictions based on the neural population’s activity during each response window. The decoding performance was evaluated using balanced accuracy.

### Response profile identification

To identify onset and sustained response for each neuron and sound, we used a non-parametric, paired Wilcoxon signed-rank test comparing baseline activity to onset or sustained activity respectively. The alpha value of the test was 0.05, implying that the proportion of neuron-pairs with significant responses has to be above 5% to exceed the maximum chance level. For offset response detection, we used two non-independent paired Wilcoxon signed-rank tests comparing offset activity to baseline and sustained activity respectively, with an alpha value of 0.05 for each test. Because the single trial measurements of baseline and sustained activity are done at a short time interval, they are not statistically independent. Therefore, the false discovery rate (FDR) is not the product of alpha values as in the case of two independent tests, but is likely higher. To be conservative, we used the alpha value of one signed-rank test (0.05, same for both tests), as an estimate of the false detection rate (FDR, i.e. chance level) for offset responses. Significant responses were classified as positive or negative based on the sign of the time-averaged baseline-substracted activity over the analysis time bin. To estimate the fraction of neurons with at least one offset response (independent of its sign), we used the Benjamin-Hochberg procedure on each neuron to correct FDR for across the 119 (dataset 1) and 112 (dataset 2) signed-rank tests performed on each sound. Therefore the chance level for the fraction of neurons responding with at least one offset is also 5%. The degrees of freedom of the signed-ranked test correspond to the number of sound repetitions: N=12 for dataset 1, N=15 for dataset 2.

### Linear decomposition of responses

In order to separate the onset, sustained and offset components, we used a linear model to approximate the response function. The linear model is a sum of three regressors (on, off and sustained) with fixed shape but whose amplitude is fit to the neural response. Since we do not know with sufficient precision the underlying biophysical mechanisms resulting in the response shape which are moreover likely to verify between neurons, we empirically designed the shape of the three regressors to resemble the data.

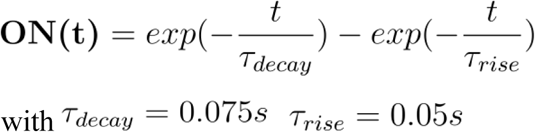

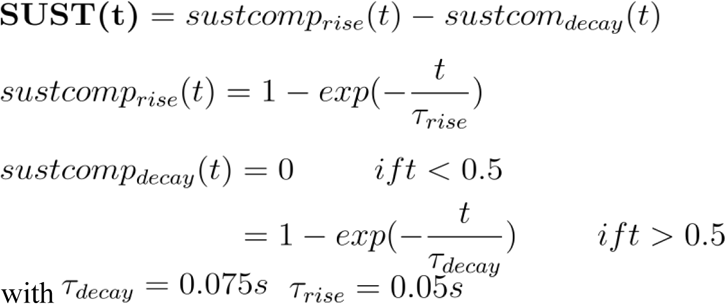

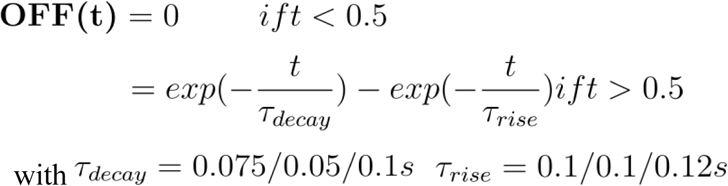

After defining three regressors, the neural response is defined by this linear equation:

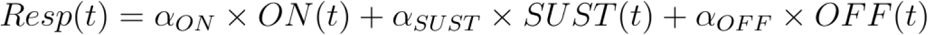

The amplitudes (αON, αSUST, αOFF) can be directly calculated (here, using Matlab’s backslash function). Given the importance of correctly fitting the offset response, we performed the fitting procedure with three different time constants and retained the best for further analysis. This fitting procedure was performed for all neurons, all sounds and every trial and the amplitudes were then used to perform classification as previously described.

## Notes

### Competing Interest Statement

The authors have declared no competing interest.

